# A New Approach to Monitoring Protein Transfer via Extracellular Vesicles

**DOI:** 10.1101/2024.12.16.628642

**Authors:** Alzbeta Chabronova, Olivia Pigden, Rosalind Jenkins, Anders Jensen, Emily Clarke, Victoria James, Mandy J Peffers

## Abstract

Extracellular vesicles (EVs) allow the exchange of protein, lipids and genetic material for communication between cells. We developed a method to track protein exchange between cells via EVs, using stable isotope labelling of amino acids in cell culture (SILAC).

We compared EV isolation methods: ultracentrifugation and size exclusion chromatography (SEC) and undertook characterisation through nanoparticle tracking analysis (NTA), then optimised the requirement (0-10%) for foetal calf serum (FCS) and EV application concentration (6.5 million-200 EVs/cell seeded). We employed heavy amino acids L-Arg-HCl and L-Lys-2HCl to label donor cells and subsequent EV proteins. Unlabelled recipient cells were treated for 12 hours with EVs, at various concentrations. Mass spectrometry proteomics was used to assess uptake of heavy labelled EVs into recipient cells.

A labelling efficiency of 62% was achieved. There was no relationship between the number of EVs applied to cells and the number of heavy proteins detected in the recipient cells. Pathway analysis indicated an inflammatory effect of applying EVs to cells, but the potential cause of this was unclear.

We identified transferred protein cargo from EVs in recipient cells. Further optimisation of this method is still necessary.

## 2. Introduction

EVs form through outward budding of the cellular membrane, encapsulating RNA, protein and lipids, facilitating communication between cells^1,2^. EV cargo is affected by the cellular environment such as in disease, leading to cellular alterations that contribute to disease pathology^2–6^. A technique to track RNA cargo exchange between cells via EVs uses 5-ethynyl uridine labelling^7^. Despite the suggested importance of transferred protein cargo over RNA exchange, (30% of the variance in an induced osteoarthritic model was explained by proteins compared to <10% from RNAs^6^), an optimised method to assessing EV protein transfer is lacking.

Currently there is little standardisation regarding isolation, detection and characterisation of EVs. There is minimal information for studies of EVs (MISEV) guidelines proposed^8–12^, however there is no gold-standard and guidelines suggest transparency when reporting rather than an optimal method. Ultracentrifugation and size exclusion chromatography (SEC) are recognised methods for EV isolation^3,6,13^, though ultracentrifugation can be harsher on the EV membrane potentially releasing EV-bound cargo^13,14^. SEC is proposed to allow greater preservation of the EV membrane, and thus the protein cargo^13^. Others suggested that ultracentrifugation resulted in increased EV purity^15^, although studies can be difficult to compare. The recent MISEV2023 guidelines propose western blotting to detect EV markers such as tetraspanins and other abundant EV proteins as validation for isolation^12^.

There is no established method to track the exchange of EV protein cargo between cells. We examined whether there was a maximal number of EVs that could be applied to cells without causing cellular stress, and whether there was a relationship between the number of transferred heavy proteins identified from the donor cells via EVs to the recipient cells and the number of heavy EVs applied. Our overarching aim was to develop a SILAC-based method to label donor cell originating proteins within EVs and to detect them using mass spectrometry in the recipient cells. Bioinformatics was used to determine potential downstream pathways affected by these EV-derived proteins. We hypothesised that using this method, we would be able to detect heavy proteins within the recipient cells indicating protein transfer from the donor cells to recipient cells via EVs.

## 3. Materials and Methods

All reagents used were from Sigma-Aldrich unless otherwise stated.

### 3.1 Ethics

We did not require ethics for our studies due to the use of cell lines.

### 3.2 Cell culture

#### 3.2.1 Standard Culture

Human SW1353 (ATCC) chondrosarcoma cells were maintained in standard culture in Dulbecco’s Modified Eagle’s Medium/Nutrient Mixture F-12 (DMEM F-12) (Gibco), (supplemented with 10% FCS (ThermoFisher) and 1% Penicillin-Streptomycin (PS) (SLS) until 80-100% confluent and trypsinised (Gibco) every 2-3 days, washing with phosphate buffered saline (PBS) (Corning/Lonza). Cells were incubated at 37 °C with 5% CO_2_.

#### 3.2.2 Stable Isotope Labelling of Amino Acids in Cell Culture (SILAC)

Cells were cultured in DMEM F-12 SILAC media (Lys and Arg-free) (ThermoFisher) supplemented with heavy arginine (L-Arg-HCl) (147.5 mg/L) (Cambridge Isotope Laboratories) and heavy lysine (L-Lys-2HCl) (91.25 mg/L) (Cambridge Isotope Laboratories)^17–19^, both sterilised using a 0.22 um filter, 10% dialyzed FCS (ThermoFisher) and 0.5% PS. To ensure appropriate incorporation of heavy labelling into the cellular proteome, cells were cultured for 10 passages.

### 3.3 Extracellular Vesicle (EV) Isolation and Characterisation

#### 3.3.1 Size Exclusion Chromatography (SEC)

Cells were washed with PBS after reaching confluence and 10% EV-free FCS (ThermoFisher) replaced FCS to supplement the media. SEC was used unless stated otherwise. We processed the media with a series of centrifugations at 300 *g* for 10 min, 2000 *g* for 10 min and then 10,000 *g* for 30 min at 4 °C to remove dead cells and debris. We then concentrated the media using VivaSpin, applying 20 mL media onto each column and centrifuging at 2000 *g* at 4 °C until the volume was approximately 0.5 mL.

EVs used in heavy labelled experiments were isolated following VivaSpin using qEV10 70nm Gen2 columns (Izon). EVs eluted in F-PBS were stored between -20 °C and -80 °C depending on time until processing.

For experiments involving application of EVs to cells, we further concentrated EVs using Amicon Ultra-15 Centrifugal Filter Unit (10 kDa) (Amicon) at 2500*g* at 4 °C to remove excess F-PBS with a final volume of approximately 2 mL (unlabelled EVs) and 1 mL (labelled EVs).

#### 3.3.2 Ultracentrifugation

For the isolation comparison experiment, we used SEC and ultracentrifugation. Cells were grown to 50-70% confluence and media harvested. Media was then processed as per Section 3.3.1. For ultracentrifugation, samples were centrifuged at 100,000 *g* for 70 min at 4 °C. Once completed, 100 μL F-PBS was pipetted into the first tube and sequentially concentrated across all tubes, resulting in all harvested EVs in one 200 μL aliquot. EVs from both isolation methods were then characterised (Section 3.3.3).

#### 3.3.3 EV Characterisation

EVs were analysed through nanoparticle tracking analysis (NTA), either using the NanoSight (Malvern Analytical) or ZetaView (Analytik). For the NanoSight, the chamber was sequentially washed with F-PBS three times. Samples were diluted until the measurement read approximately 10^8^ EVs and then scaled to the original concentration. Analysis was conducted using NanoSight NTA 3.4. Unless otherwise stated, EV concentrations were reported using ZetaView measurements.

### 3.4 Experimental Culture

For all experiments, cells were seeded at 30,000 cells/cm^2^ a day prior to treatment.

#### 3.4.1 Application of Unlabelled EVs to Cells

EV tolerance experiments assessed cell stress using unlabelled EVs. Cells were cultured in standard conditions until replacing the media once cells were 100% confluent with DMEM-F12 GlutaMAX (Gibco) supplemented with 0.1% FCS and 1% PS. To collect media containing EVs, cells were cultured for approximately 4-6 days, refreshing media every 2-3 days. Media was stored at –80 °C until processing at 300 *g* for 10 min, 2000 *g* for 10 min, then 10,000 *g* for 30 min at 4 °C. EVs were isolated using SEC (Section 3.3.1).

To apply the EVs to cells, cells were seeded in 24-well plates in DMEM F-12 with 10% EV-free FCS and 1% PS. The next day, EVs were applied to cells as detailed in **Table 1** in triplicate technical replicates (n=3) in a serial dilution following filter sterilisation with a 0.22 μm filter. Cells were then visually assessed after 24 hours and 48 hours.

**Table 1.** Concentrations of unlabelled EVs applied to unlabelled cells in serial dilution to assess the cells’ tolerance to EVs.

| *Code* | *EVs/cell seeded (ZetaView)* |
| --- | --- |
| *A* | 200 |
| *B* | 400 |
| *C* | 800 |
| *D* | 1600 |
| *E* | 3200 |
| *F* | 6400 |
| *G* | 12800 |
| *H* | 25600 |
| *I* | 51200 |
| *J* | 102400 |
| *K* | 204800 |
| *L* | 409600 |
| *M* | 819200 |
| *N* | 1638400 |
| *O* | 3276800 |
| *P* | 6553600 |

#### 3.4.2 FCS Supplementation Requirements

To assess FCS requirements of the cells, cells were seeded at 30,000 cells/cm^2^ in 12-well plates. Media was changed the following day with either 0%, 0.1%, 10% FCS or EV-free FCS and a seventh condition where cells were not refreshed (control 1), 10% FCS refreshment was considered control 2. Visual assessment of confluence at 3, 5.5 and 24 hours was used as a proxy for cell viability/stress.

#### 3.4.3 Cell Viability Assessment

To assess cell viability quantitatively, we used AlamarBlue^TM^ Cell Viability Reagent (Invitrogen). Cells were seeded cells in 96-well plates in proliferation media, then treatment media applied the following day. Treatment media consisted of EV concentration-A, -H and -P (**Table 1**) in either no FCS or 10% EV-free FCS, 10% FCS with no EVs, 10% EV free FCS with no EVs, or no FCS with no EVs and 40% Dimethyl Sulfoxide (DMSO) as a negative control. Alamar blue was added following manufacturer instructions and colour intensity measured using fluorescence at excitation 560 nm and emission 590 nm, normalised to background colour intensity after adjusting the gain for well volume. Cells were incubated for 2 hours following a 4-hour incubation with the treatment media (total 6 hours of treatment).

#### 3.4.4 Application of Heavy EVs to Recipient Cells

Cells were seeded at 30,000 cells/cm^2^ in 6-well plates in triplicate for each concentration (**Table 2**) in standard unlabelled proliferation media. The following day, cells were washed with PBS and refreshed with DMEM F-12 supplemented with 10% EV-free FCS, 1% PS and the EV serial dilutions. The EV-containing treatment media was incubated on the cells for 12 hours before harvesting the cells, as detailed in Section 3.3.1.

**Figure 1.**
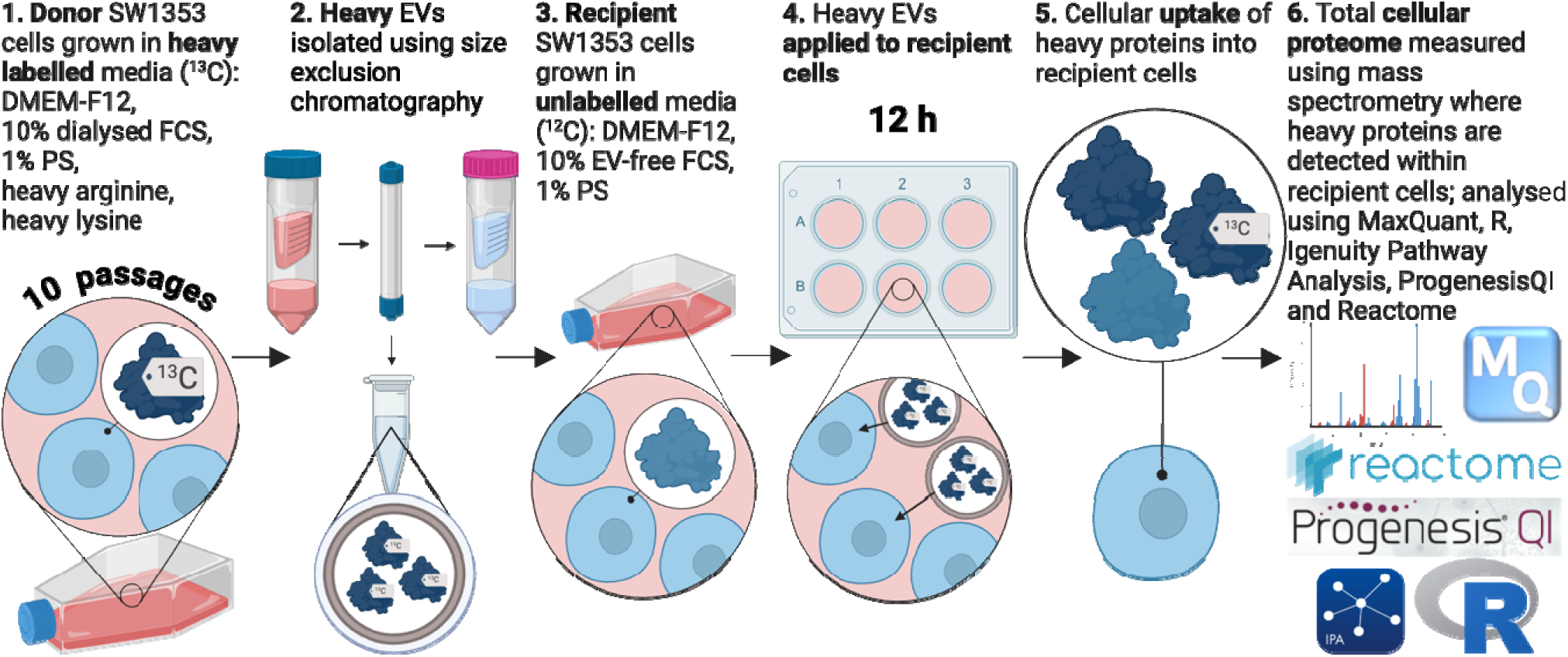
Heavy-labelled EV experiment workflow. SW1353 cells grown for 10 passages to achieve complete heavy-label incorporation of all proteins within the donor cells. We then isolated EVs in filtered PBS from this using SEC using a 100 kDa VivaSpin, qEV10 Izon column followed by a 10 kDa Amicon filter to concentrate the sample. These heavy-labelled EVs were then applied to unlabelled recipient cells in regular proliferation media with 10% EV-free FCS for 12 hours (n=3 for each concentration condition ranging from 6 million (A) to 600 (E) EVs per seeded cell). The recipient cellular proteome was then analysed using mass spectrometry to detect the transferred heavy-labelled donor cell EV originating proteins.

**Table 2.** Concentrations of heavy-labelled EVs applied to unlabelled cells in serial dilution.

| Concentration Code | Raw number of EVs (particles/mL) (ZetaView) | Number of EVs/cell seeded (30,000 cells/cm<sup>2</sup>) |
| --- | --- | --- |
| A | 1.7x10<sup>12</sup> | 6x10<sup>6</sup> |
| B | 1.7x10<sup>11</sup> | 6x10<sup>5</sup> |
| C | 1.7x10<sup>10</sup> | 6x10<sup>4</sup> |
| D | 1.7x10<sup>9</sup> | 6x10<sup>3</sup> |
| E | 1.7x10<sup>8</sup> | 6x10<sup>2</sup> |
| No EV (control) | 0 | 0 |

### 3.5 Proteomics

#### 3.5.1 Protein Extraction

Cells treated with EVs in a 6-well plate (n=3 for each concentration) were washed with PBS twice and scraped on ice into a 200 μL solution of 25 mM Ammonium Bicarbonate (AmBic) in LC-MS grade water (ThermoFisher), 0.15% benzoase nuclease (Millipore) and a proteinase inhibitor tablet diluted to 1x concentration per well. To lyse the cells and extract the protein, cells were sonicated at 10 microns fourteen times for 1 s on and a 1 s rest between cycles. This wa then centrifuged at 13,000 rpm for 10 min at 4 °C. Supernatant was then removed. Cell lysate protein concentration was determined using Pierce BCA Protein Assay Kit (Thermo Scientific) following manufacturer guidance.

#### 3.5.2 Protein Digestion

##### 3.5.2.1 Cellular Protein Digestion

RapiGest SF Surfactant (Waters) was diluted in 100 μL of 25 mM AmBic dissolved in LC-MS grade water to make a 1% (w/v) solution. 10 μL aliquots of this were added to each sample containing 80 ug of protein. This was incubated at 80 °C for 10 min. 10 μL of 11mg/mL DL-Dithiothreitol (DTT) solution (Merck) was then added and incubated at 60 °C for 10 min. In the dark, 10 μL of 46 mg/mL Iodoacetamide (IAA) was then added for 30 min. Subsequently, 9.4 μL of DTT was then added. 10 μL of Trypsin/Lys-C Mix (Promega) dissolved in 100 uL 25 mM AmBic in LC-MS grade water and incubated at 37 °C overnight with a top up of 10 μL trypsin after 2 hours. Samples were acidified with trifluoroacetic acid (TFA), incubated at 37 °C for 45 min followed by centrifugation at 13,000 *g* for 15 min at 4 °C. A 10 μL aliquot of this was taken along with a 10 μL aliquot pre-digest and run on a gel to ensure full protein digestion. For the protein gel, pre- and post-digest aliquots were run using a NuPAGE 4 to 12% Bis-Tris gel (Invitrogen). Samples were combined with a solution of 2 x SDS sample buffer (ThermoFisher) with 5 mM DTT. Samples were then boiled at 95 °C for 10 min and placed on ice before applying to the gel at 20 μL per sample. One lane contained 10 μL Novex Sharp Pre-Stained Protein Standard ladder (Invitrogen). Gels were run at 200 V for 20 min. The gel was then washed and placed in Colloidal blue (Invitrogen) following manufacturer guidance.

##### 3.5.2.2 EV Protein Digestion

For the EV proteomics, we used the same workflow as Section 3.5.2.1 however the sample (120 μL EVs (circa. 2.1×10^12^ (ZetaView)) suspended in F-PBS) was suspended in 60 μL 3x concentrated urea lysis buffer (6M urea, 1M AmBic in MQ, 0.5% deoxycholate) before sonication to assist membrane lysis. Due to the low EV protein content of EVs, we diluted the DTT and IAA to a 5x dilution. To ensure complete protein digestion, samples were run on a gel using a 10 μL aliquot both pre and post-digest. 3 μL NuPAGE LDS sample buffer (Invitrogen) (92 μL stock diluted in 8 μL 2-Mercaptoethanol) was added to the 10 μL samples, then heated at 95 °C for 10 min. This was then run on a NuPAGE 4 to 12% Bis-Tris gel with 1 x NuPAGE MES running buffer (Invitrogen). Samples were then run alongside 5 μL of Novex Sharp Pre-Stained Protein Standard ladder at 180 V for 30 min. Gels were visualised following Colloidal blue staining kit (Invitrogen) following manufacturer guidance.

#### 3.5.3 Label-Free liquid chromatography-tandem mass spectrometry (LC-MS/MS)

Approximately 750 ng of each digested cell lysate was analysed using LC-MS/MS on a 2-hour gradient with a 10x diluted final sample (untreated cell lysate) as described by Johnson *et al.*^20^. Heavy EV proteomics were run with the same settings, with a sample containing 2.1×10^12^ EVs.

#### 3.5.4 Peptide and SILAC label identification and quantification

MaxQuant was used to identify any heavy-labelled proteins as described by Cox *et al*.^21^. For recipient cell proteomics, proteins with a heavy/light ratio were identified as heavy labelled and for the EV proteomics, proteins with heavy but not light and a mix of light and heavy signal were identified as heavy. To calculate a labelling efficiency for the cell lysates, we ran evenly split light/heavy mixes (n=3) and counted the number of heavy proteins divided by the total number of proteins.

For protein identification, we used a local Mascot server searching ‘UniHuman’ database with fixed modifications at ‘Carbamidomethyl (C)’ and variable modifications at ‘Oxidation (M)’, ‘Label: 13C(6) (K)’ and ‘Label:13C(6) (R)’ and peptide charge at ‘2+ and 3+’ with a peptide mass tolerance at 10.0 ppm and 0.01 Da fragment mass tolerance, and one missed cleavages allowed. For cell lysates, label-free protein quantification was undertaken using Progenesis QI (Waters)^20^. The Progenesis normalised values for each comparison control (n=3) versus treatment group (A-E) (n=3) were then used to assess differential abundance, comparing three methods (Section 3.6.2).

### 3.6 Data Analysis

#### 3.6.1 Statistical Analysis

Cell viability assessment from the Alamar blue assay was analysed using R Studio (version 4.3.2 -- “Eye Holes”) with the following packages: “nparcomp”, “ggplot2”, “dplyr”, “stringr” and “RColorBrewer”, first testing normality using a Shapiro-wilk test, then a Kruskal-Wallis test followed by a Tukey multiple comparisons test.

To test for a relationship between the number of heavy proteins identified and number of EVs applied to cells, the MaxQuant number of heavy proteins and number of EVs applied to cells (A-E) as detailed in **Table 2** were checked for normality using a Shapiro-wilk test and then a Spearman’s correlation in R Studio.

#### 3.6.2 Proteomic Analysis

Due to the low labelling efficiency, we also assessed differential abundance from the total proteome as well as the heavy labelled proteins to assess the global effects of EV application to cells. We evaluated three different methods for analysing the total proteome, all at threshold cut-offs of 1.3-FC and p or q-value <0.05.

The Progenesis normalised values (normalised to each other based on the chromatograms produced from mass spectrometry) for each comparison control (n=3) vs treatment group (A-E) (n=3) were used to calculate a FC using treatment divided by control values and then log2 transformed, the q-value was also log10 transformed. This analysis formed the first Progenesis method.

The second method involved running the normalised values from Progenesis and gene names but ignoring the statistical analysis and performing this in R Studio instead. For this, the following packages were used: “RColorBrewer”, “ggrepel”, “tidyverse”, “readxl”, “SummarizedExperiment”, “NormalyzerDE”, “tibble”, “tidyr” and “readr”. This allowed quantile normalisation of the *limma* differential expression data for statistical analysis and producting a p-adj value with Benjamini-Hochberg correction.

The final method involved the same packages as the second method as well as: “dplyr”, “stringr”, “reshape2”, “ggplot2”, “VIM”, “missMethods”, “pcaMethods”, “missForest”, “ggpubr”, “janitor”, “ggvenn”. This method removed values that were only contained within either the treatment or control group. To classify this, as we had three replicates for each concentration, if only one of the three replicates contained a value, this was assumed a false positive and the protein was classed as not present in that group. If there was a protein in which two measurements were present, but a missing value was in the third replicate, the third value was imputed. First the data was normalised as per the previous method with quantile normalisation. Proteins were then removed and assigned to a separate list of proteins unique to either treatment, or control to be displayed in a Venn diagram.

To assess the best method for imputing the missing values, k-Nearest Neighbors, Local Least Squares, random forest and Bayesian Principle Component Analysis (BPCA) were assessed using a machine learning loop (n=50) where the method with the lowest and least variation of the sum of squared errors (BPCA) was chosen to impute these missing values (**Table 3**; differential abundance was filtered to 1.3-FC and using either q-value for Progenesis analysis, or in both methods involving R analysis, p-adj value with Benjamini-Hochberg correction at < 0.05).

**Table 3.** Mean percentage confluence based on visual assessment of SW1353 cells incubated for 24 and 48 hours at varying concentrations of EVs.

| *EV concentration*<br>*treatment code* | *EVs/cell seeded*<br>*(ZetaView)* | *24 hrs* | *48 hrs* |
| --- | --- | --- | --- |
| *A* | 200 | 97 | 100 |
| *B* | 400 | 97 | 100 |
| *C* | 800 | 97 | 100 |
| *D* | 1600 | 95 | 100 |
| *E* | 3200 | 67 | 75 |
| *F* | 6400 | 55 | 67 |
| *G* | 12800 | 73 | 88 |
| *H* | 25600 | 63 | 72 |
| *I* | 51200 | 90 | 97 |
| *J* | 102400 | 77 | 93 |
| *K* | 204800 | 85 | 97 |
| *L* | 409600 | 97 | 98 |
| *M* | 819200 | 82 | 97 |
| *N* | 1638400 | 72 | 92 |
| *O* | 3276800 | 65 | 82 |
| *P* | 6553600 | 73 | 88 |

#### 3.6.3 Pathway analysis: Reactome DB and Ingenuity Pathway Analysis (IPA)

Downstream pathway analysis of heavy proteins was undertaken using Reactome DB (https://reactome.org).

For cell proteomic pathway analysis, we used Ingenuity Pathway Analysis (IPA) (Ingenuity Systems) to determine canonical pathways and upstream regulators of the proteins using the Progenesis data following imputation in R (as described above) at thresholds of p-adj < 0.05 and 2 FC using the user defined dataset as background (all proteins identified). A Right-tailed Fisher’s exact test was used to calculate the p-values in IPA.

#### 3.6.4 FunRich EV protein enrichment analysis

FunRich version 3.1.4^22,23^ was used to allow identification of proteins overlapping with the Vesiclepedia^24^ within the software and user defined datasets. We compared proteins between this dataset, heavy EV proteome and heavy proteins identified in cell lysates from all concentrations (A-E) overlapping in two or more replicates (n=3) after removing false positives identified in untreated cells.

**Data Availability Statement:** Proteomics data are available via ProteomeXchange with identifier PXDXXXX.

## 4. Results

### 4.3 Applying unlabelled EVs to cells did not cause stress using visual assessment

The primary assessment of cell stress caused by application of EVs was performed using visual assessment of confluence (**Table 3**; percentage confluence was estimated by visual assessment of three wells for each condition, then calculating the mean). Results showed no direct relationship between the number of EVs applied and cell stress as determined by visual assessment of confluence as a proxy for stress (n=3).

Next, we used Alamar blue to assess cell viability following the application of varying EV concentrations in either 0% or 10% FCS. We found a significant difference in cell viability following the application of EVs in 10% EV-free FCS (p=0.0003625). This was significant for comparisons between DMSO and each treatment but not between treatments ( DMSO vs 10% EV-free FCS, p=0.0000000, DMSO vs 10% EV-free FCS supplemented with EVs at concentration-A, p=0.0000000, DMSO vs 10% EV-free FCS supplemented with EVs at concentration-H, p=0.0000000, DMSO vs 10% EV-free FCS supplemented with EVs at concentration-P, p=0.0000000) (**Figure 2**). Similarly, we found a significant difference when EVs were in 0% FCS (p=0.0002121), again this was between the DMSO negative control and each treatment group (**Figure 3**).

**Figure 2.**
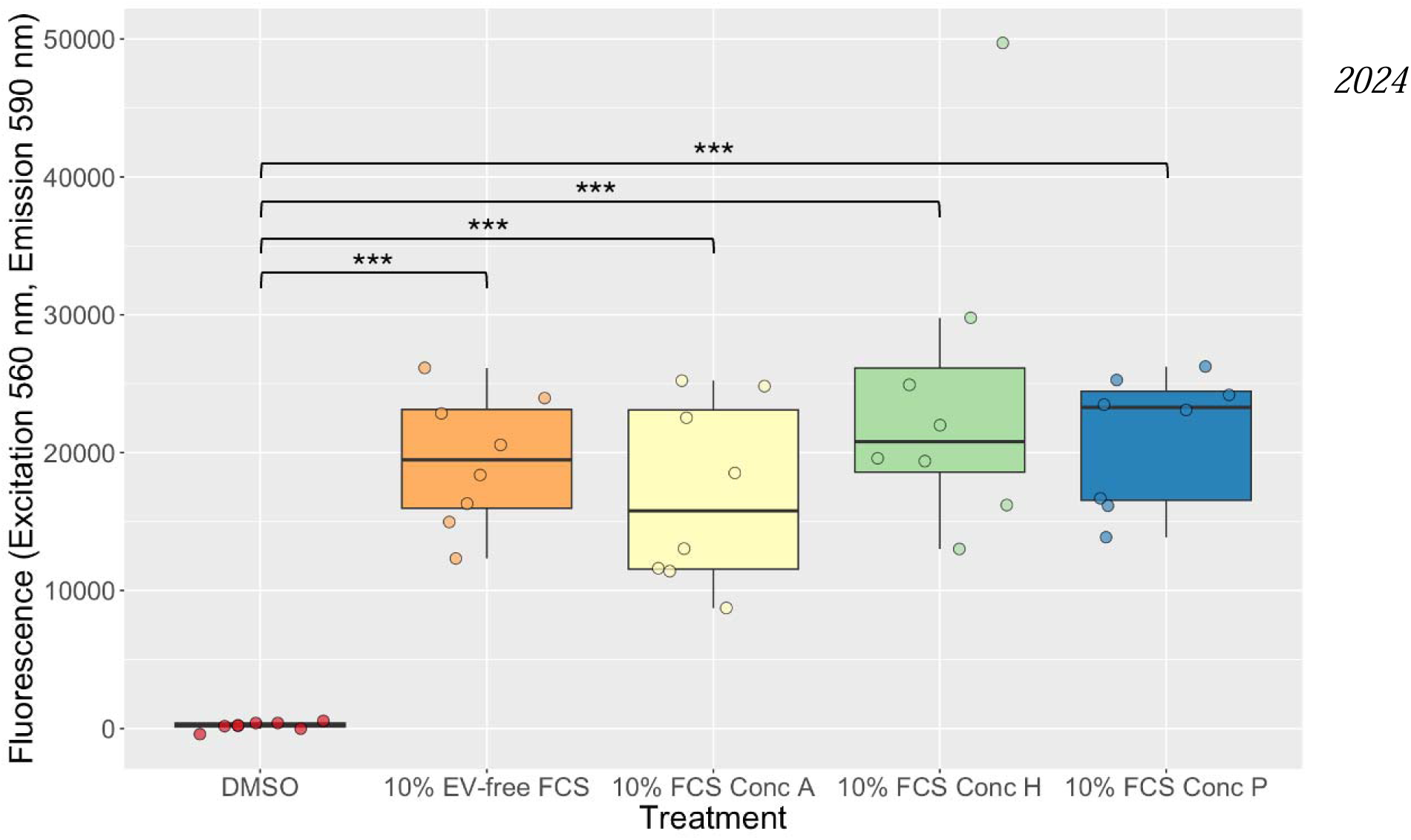
Alamar blue cell viability assessment of SW1353 cells in 10% EV-free FCS following EV application for 6 hours (n=8). SW1353 cells cultured in media supplemented with 10% FCS and varying concentrations of EVs. SW1353 cells were seeded in 10% FCS proliferation media one day prior to washing with PBS and application of treatment media for 6 hours in total with Alamar blue reagent added for the final 2 hours. Treatment medias were either 40% DMSO in 0% FCS (n=8) as a negative control, 10% EV-free FCS with no EVs (n=8), 10% EV-free FCS with concentration-A = 200 EVs per seeded cell (n=8), 10% EV-free FCS with concentration-H = 25600 EVs per seeded cell (n=8) or 10% EV-free FCS with concentration-P = 6553600 EVs per seeded cell (n=8). Fluorescence, adjusted gain for well (1499), was measured at excitation at 560 nm and emission at 590 nm, with background media colour removed from control media of 0% or 10% EV-free media. Data was plotted in R using packages “ggplot2”, “dplyr”, “stringr”, “RColorBrewer”, Kruskal-Wallis and post-hoc Tukey tests were used to analyse the data (*** p<0.001).

**Figure 3.**
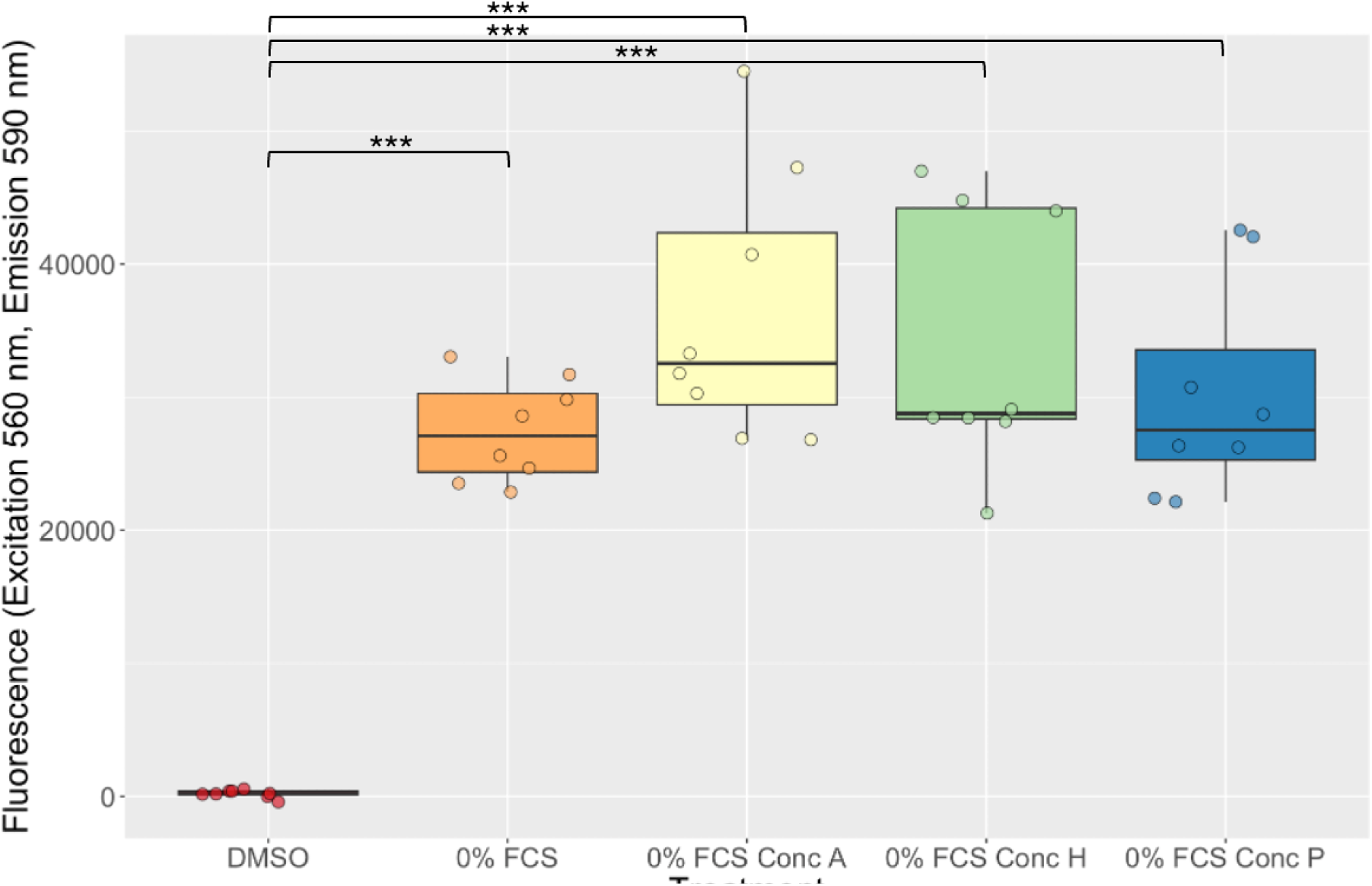
Alamar blue cell viability assessment of SW1353 cells in 0% EV-free FCS with following EV application for 6 hours (n=8). SW1353 cells cultured in media supplemented with 0% FCS and varying concentrations of EVs. SW1353 cells were seeded in 10% FCS proliferation media one day prior to washing with PBS and application of treatment media for 6 hours in total with Alamar blue reagent added for the final 2 hours. Treatment medias were either 40% DMSO (dimethylsulphoxide) in 0% FCS (n=8) as a negative control, 0% FCS with no EVs (n=8), 0% FCS with concentration-A = 200 EVs per seeded cell (n=8), 0% FCS with concentration-H = 25600 EVs per seeded cell (n=8) or 0% FCS with concentration-P = 6553600 EVs per seeded cell (n=8). Fluorescence, adjusted gain for well (1499), was measured at excitation at 560 nm and emission at 590 nm, with background media colour removed from control media of 0% FCS media. Data was plotted in R using packages “ggplot2”, “dplyr”, “stringr”, “RColorBrewer”, Kruskal-Wallis and post-hoc Tukey tests were used to analyse the data (*** p<0.001).

### 4.4 Heavy proteins originating from the donor cell EVs were detected within recipient cells

Following labelling of SW1353 cells, labelling efficiency reached 62% (n=3). Thus, EVs were not fully labelled. It was therefore likely that we missed unlabelled proteins that have been transferred from the donor EVs to the recipient. Despite this, we were able to detect heavy proteins within the recipient cells (**Figure 4**). However, we also identified heavy proteins within the untreated control cells despite them not being treated with heavy EVs indicating false positives. Therefore, we removed heavy proteins found within two or more of the three replicates.

**Figure 4.**
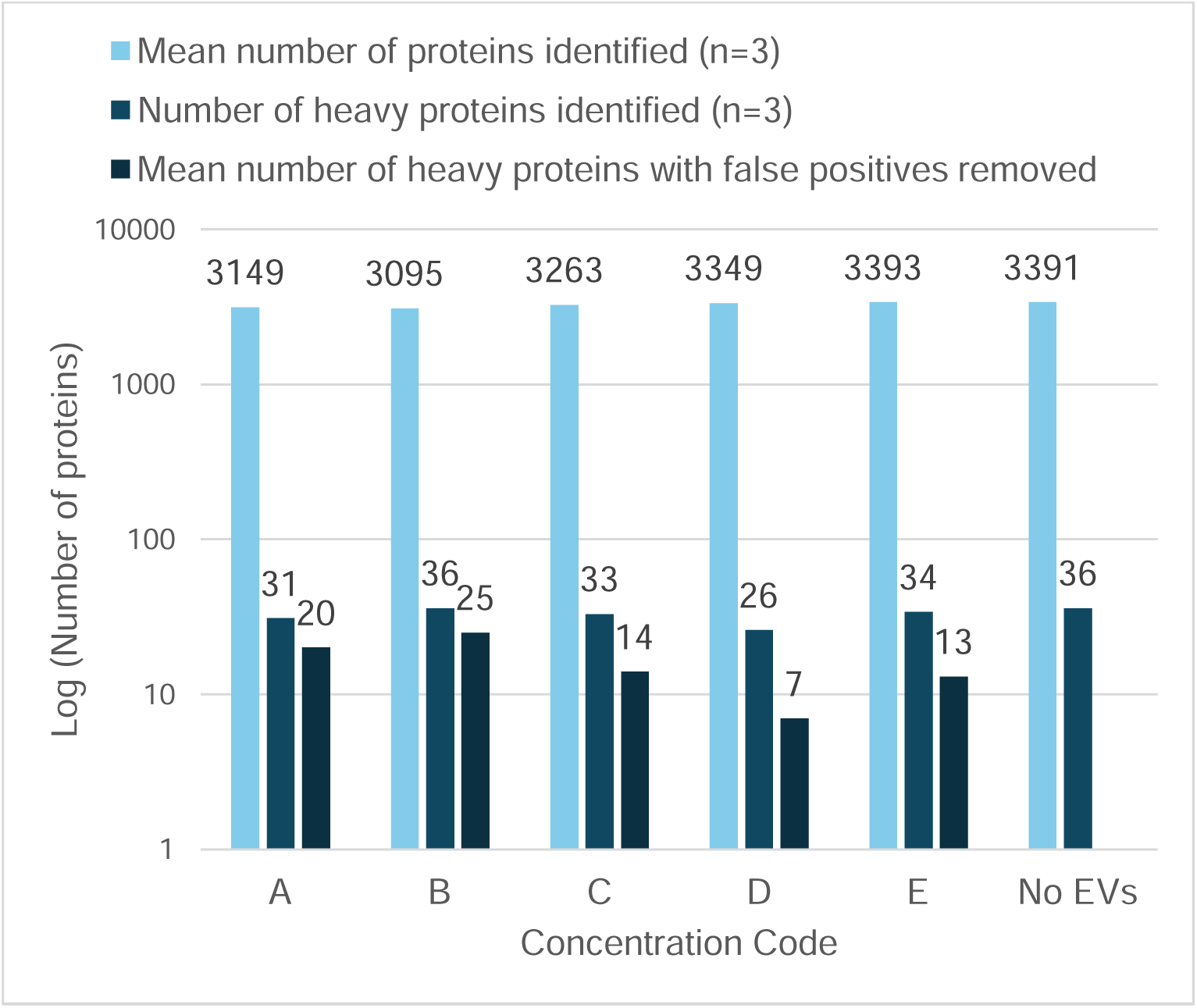
Number of heavy proteins identified within the recipient cells using MaxQuant data analysis (n=18). Heavy proteins identified appeared in two or more replicates on a logarithmic scale. This list was then compared with the heavy proteins detected in the untreated control samples to remove assumed false positives. Concentration codes are from **Table 2**.

### 4.5 We detected 22 heavy proteins originating from donor cell EVs within recipient cells

To determine if the heavy proteins that we detected originated from the applied heavy EVs, we analysed the proteome of a sample of the heavy-labelled EVs using mass spectrometry. We identified 22 heavy proteins that overlapped with the recipient cell-originating heavy proteins.

We identified 517 proteins within the heavy EV proteome that were not associated with EVs (**Figure 5**). All heavy proteins identified within the recipient cell lysates (all concentrations (A-E)) were identified within EVs previously.

**Figure 5.**
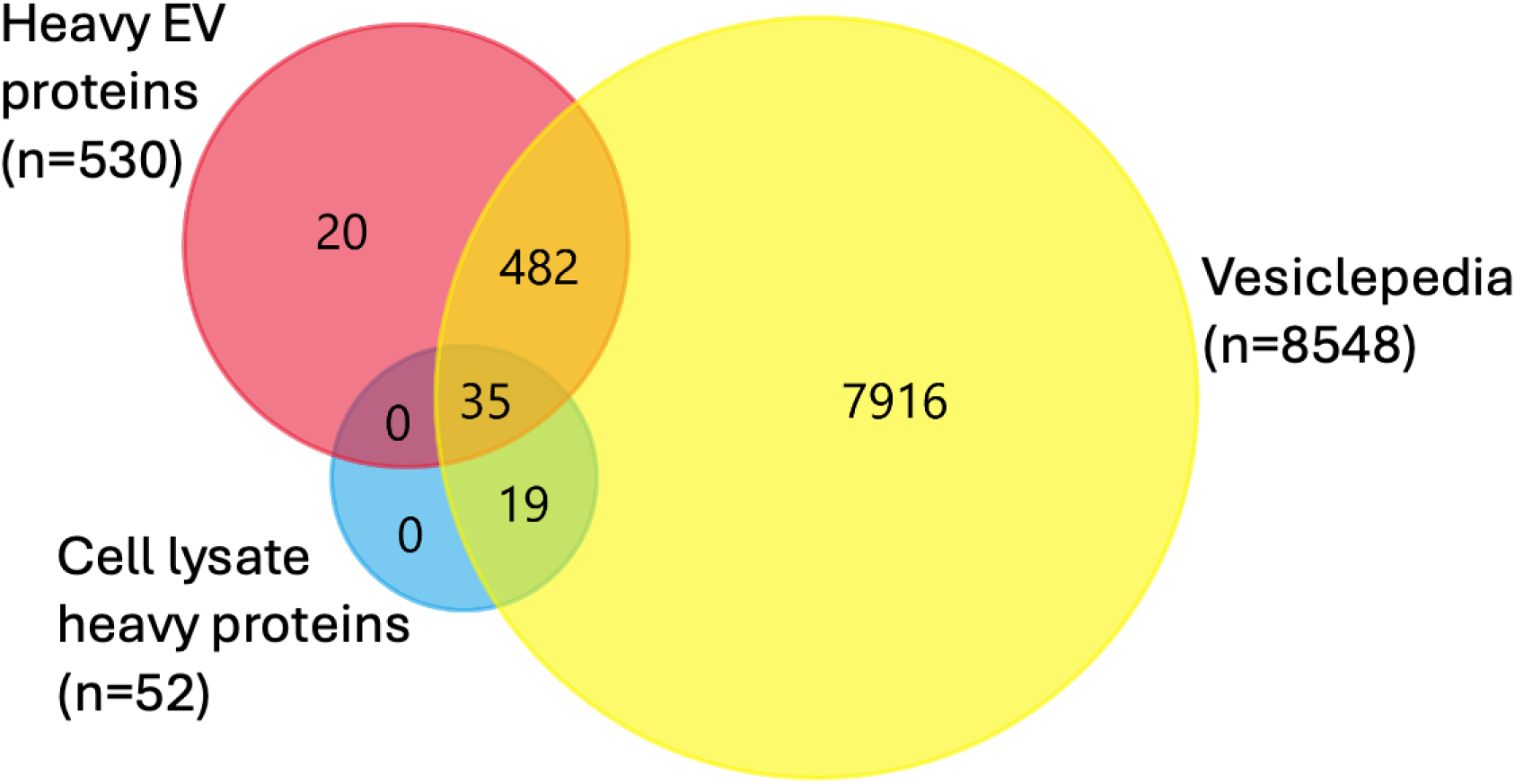
Overlapping known EV proteins with current identified heavy-labelled proteins. Proteins identified in EVs within Vesiclepedia^24^ as part of FunRich analysis^22,23^ were used to allow identification of heavy EV proteins (n=530) and heavy proteins identified in cell lysates from all concentrations (A-E) overlapping in two or more replicates (n=3) after removing false positives identified in the untreated cells (n=52).

### 4.6 Pathways of the 22 overlapping proteins mapped to cell communication, protein processing and stress

The role of the 22 overlapping proteins was investigated using Reactome DB. The top five significant pathways were associated with inflammation (**Table 4**; heavy proteins appeared in two or more of the replicates. This list was then compared with the heavy proteins detected in the untreated control samples). Other pathways of note included lysosome vesicle biogenesis (p=9.84×10^-5^), cellular responses to stress (p=1.76×10^-4^), cellular responses to stimuli (p=2.01×10^-4^), protein methylation (p=7.21×10^-4^), vesicle-mediated transport (p=1.24×10^-3^), membrane trafficking (p=2.16×10^-3^), cell-cell communication (p=4.69×10^-3^), protein folding (p=1.8710^-2^), cell-extracellular matrix interactions (p=3.83×10^-2^), metabolism of proteins (p=4.22×10^-2^), clathrin-mediated endocytosis (p=4.32×10^-2^).

**Table 4.** Top Reactome DB pathways associated with heavy proteins identified following MaxQuant analysis in all concentrations (n=15).

| Pathway | P-value |
| --- | --- |
| Gene and protein expression by JAK-STAT signalling after interleukin-12 stimulation | $3.33 \times 10^{-16}$ |
| Interleukin-12 signalling | $1.22 \times 10^{-15}$ |
| Interleukin-12 family signalling | $4.55 \times 10^{-15}$ |
| Immune System | $1.45 \times 10^{-10}$ |
| Signalling by interleukins | $3.31 \times 10^{-9}$ |

### 4.7 Cells treated with concentration-A and -B had differentially abundant (DA) heavy proteins

To determine the effects of applying the heavy EVs to cells, we assessed the cellular proteome. Additionally, we explored different methods for assessing DA proteins using Progenesis and R for concentration-A proteins. We found that the most stringent method for assessing DA proteins used a combination of Progenesis and imputing missing values in R. Therefore, we employed this method for the remaining EV treatment concentrations. To perform this analysis and calculate FC, we removed proteins unique to the control or treatment group within each comparison analysis (**Figure 5)**.

We then analysed these unique proteins collectively, classing as unique to control (n=5) (untreated unlabelled cells) or unique to treatment (n=5) (unlabelled cells treated with heavy labelled EVs), using Reactome DB to identify their roles. 33 proteins were identified as unique to treatment and 26 unique to control. Pathways of significance unique to treatment included pyroptosis (p=3.35×10^-3^), interleukin-37 signalling (p=3.10^-3^), metabolism of RNA (p=4.42×10^-3^), FOXO-mediated transcription of oxidative stress, metabolic and neuronal genes (p=6.79E-3), defective pyroptosis (p=7.33×10^-3^), PTEN regulation (p=8.73×10^-3^), death receptor signalling (p=9.16×10^-3^), epigenetic regulation of gene expression (p=1×10^-2^), regulated necrosis (p=1.57×10^-2^), release of apoptotic factors from the mitochondria (p=1.98E-2), diseases of programmed cell death (p=2.47×10^-2^), FOXO-mediated transcription (p=3.11×10^-2^), class I peroxisomal membrane protein import (p=4.89z10^-2^), post-translational protein modification (p=4.95×10^-2^). Other pathways involved associates with the unique to control proteins include TP53 regulates transcription of cell death genes (p=1.34×10^-2^), suppression of apoptosis (p=2.92×10^-2^), unfolded protein response (p=4.32×10^-2^), TP53 regulates transcription of genes involved in G2 cell cycle arrest (p=4.32×10^-2^).

In our differential abundance analysis, we found 262 proteins increased following treatment with concentration-A, 136 in concentration-B, 0 in concentration-C, 0 in concentration-D and 1 in concentration-E, and 263 proteins reduced in concentration-A, 130 in concentration-B, 0 in concentration-C, 0 in concentration-D and 1 in concentration-E at a 1.3 FC and p-adj with Benjamini-Hochberg correction <0.05 (**Figure 6**).

**Figure 5.**
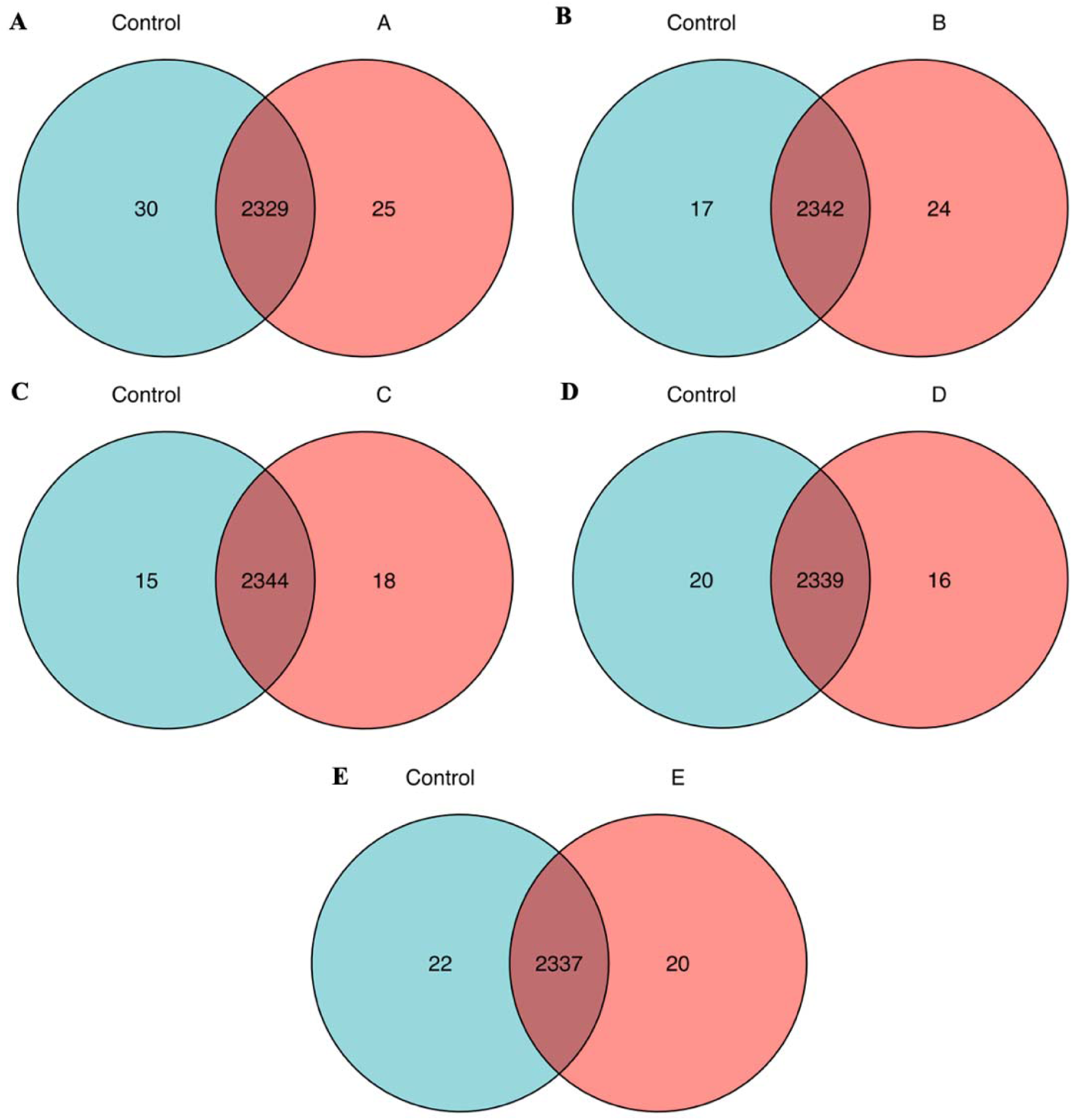
Venn diagrams of proteins uniquely identified following Progenesis analysis. Control (n=3) (light blue) or treatment groups (red) for (A) concentration-A (n=3), (B) concentration-B (n=3), (C) concentration-C (n=3), (D) concentration-D (n=3), (E) concentration-E (n=3) prior to imputation of missing values. Proteins were considered unique if they were not identified in one group, or if only present in one of three replicates and considered as false positives. Data was plotted in R using packages “RColorBrewer”, “ggrepel”, “tidyverse”, “readxl”, “SummarizedExperiment”, “NormalyzerDE”, “tibble”, “tidyr”, “dplyr”, “stringr”, “reshape2”, “ggplot2”, “readr”, “ggpubr” and “ggvenn”.

**Figure 6.**
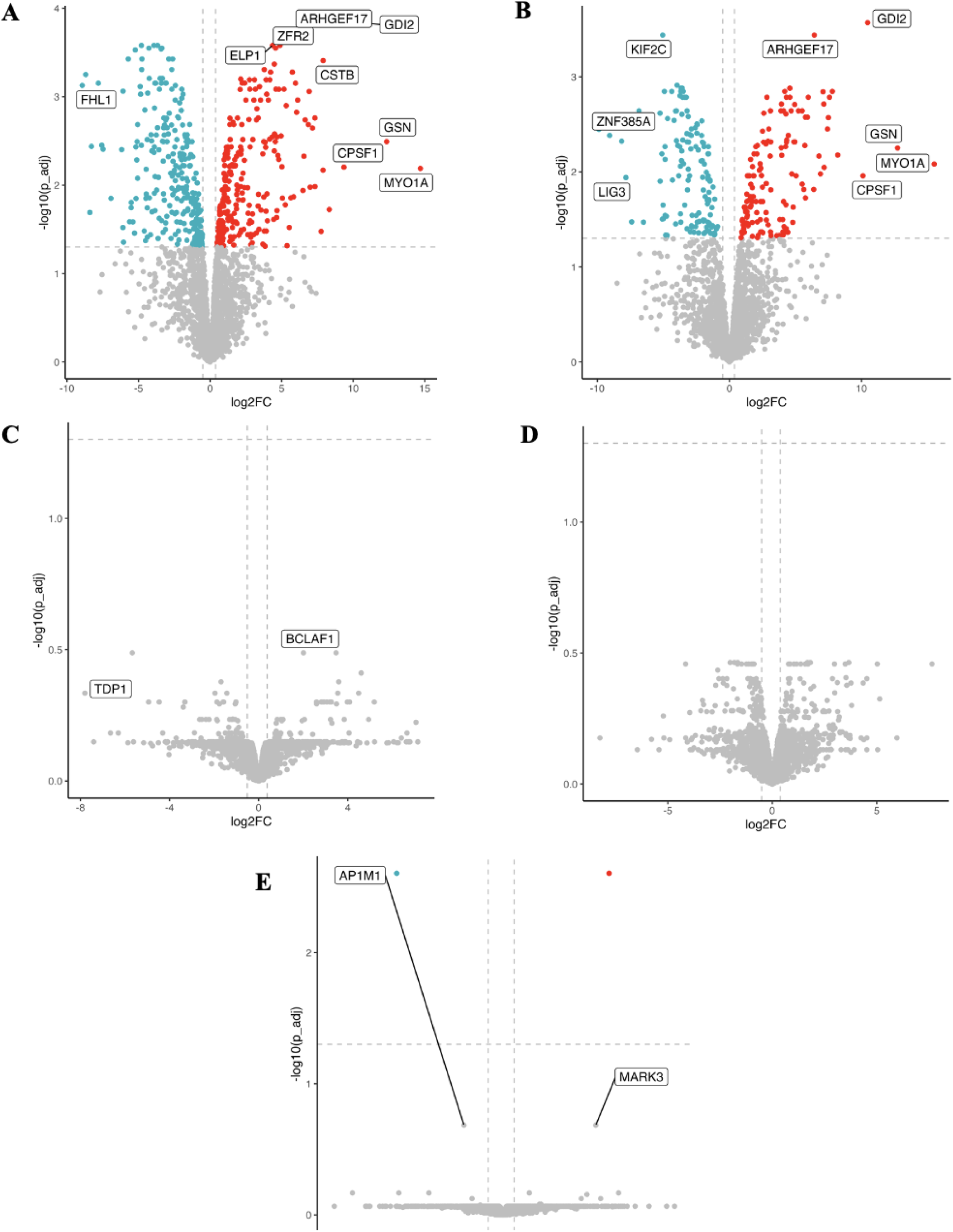
Volcano plots of DA proteins within the total cellular proteome following addition of heavy EVs. (A) concentration-A (n=3), (B) concentration-B (n=3), (C) concentration-C (n=3), (D) concentration-D (n=3), (E) concentration-E (n=3) vs no EV treatment control (n=3). Performed at cut-offs of 1.3-FC and p-adj < 0.05. Benjamini-Hochberg correction, using quantile normalisation with Bayesian PCA imputation on data where one of the three replicates was missing (n=3). There were (A) 262, (B) 136, (C) 0, (D) 0, (E) 1 protein increased (red) and (A) 263, (B) 130, (C) 0, (D) 0, (E) 1 protein reduced (light blue). Data was plotted in R using packages “RColorBrewer”, “ggrepel”, “tidyverse”, “readxl”, “SummarizedExperiment”, “NormalyzerDE”, “tibble”, “tidyr” and “readr” “dplyr”, “stringr”, “reshape2”, “ggplot2”, “VIM”, “missMethods”, “pcaMethods”, “missForest”, “ggpubr”, “janitor”.

### 4.8 Following EV treatment of cells, pathways from DA proteins mapped to EIF3E, NR3C1, SLC40A1, cell cycle and protein processing

We used IPA to investigate the upstream regulators and top canonical pathways associated with proteins identified following imputing missing values. We set cut-offs of 2-FC, and 0.05 p-value adjusted for Benjamini-Hochberg correction and found 406 DA proteins, 214 reduced and 192 increased concentration-A and 259 DA proteins, 128 reduced and 131 increased in concentration-B. In upstream regulators of the proteins found within the dataset following treatment with concentration-A, EIF3E (z=-2.449, p=3×10^-3^) was predicted as significantly inhibited. For concentration-B, SLC40A1 (z=2.138, p=2.04×10^-2^) and NR3C1 (z=2.741, p=2.34×10^-2^) were predicted as significantly activated (**Figure 7**).

**Figure 7.**
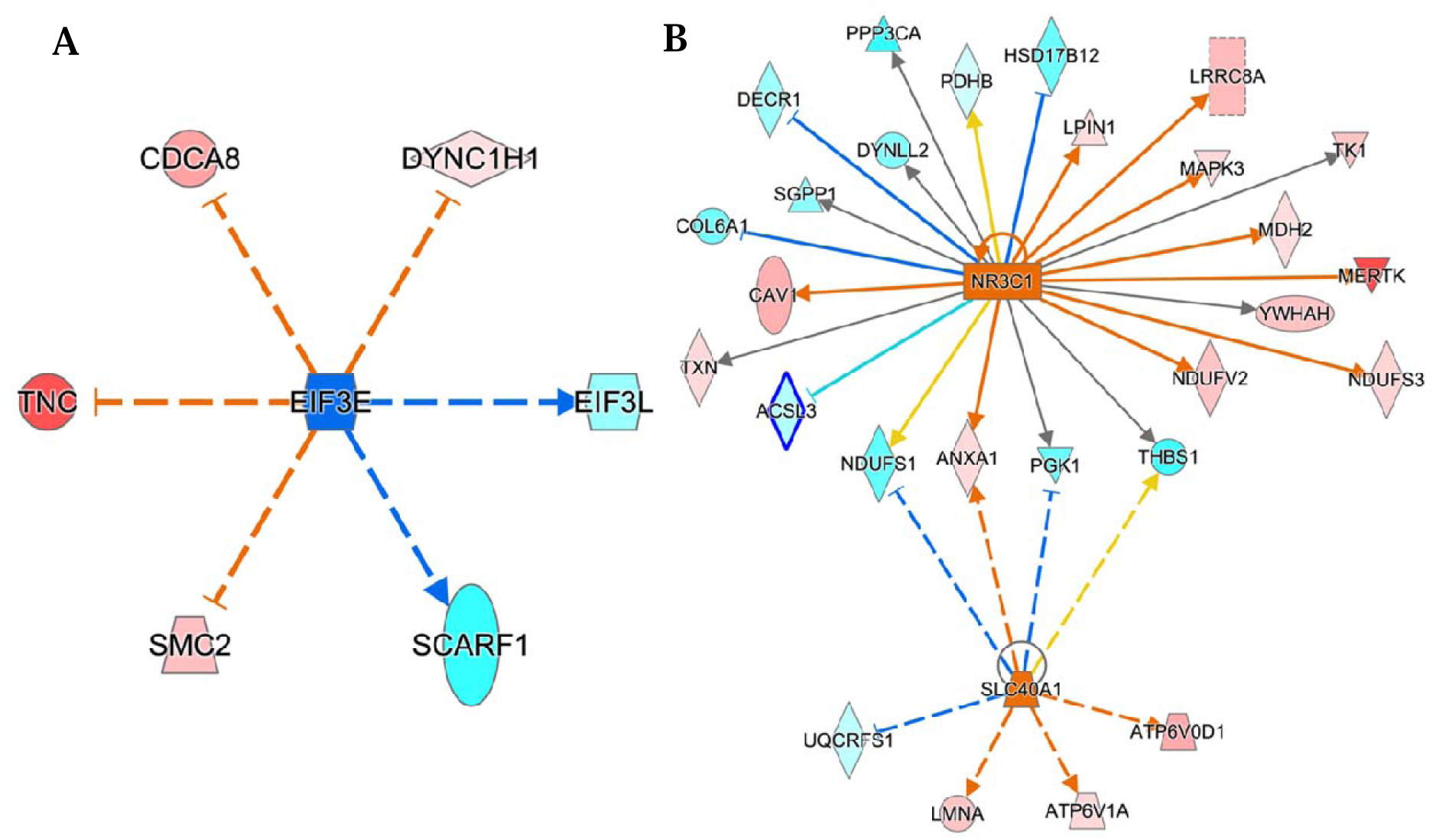
Proteins predicted to be activated (orange) or inhibited (dark blue) in IPA from proteins at p<0.05 and 2-FC for (A) A (n=3) and (B) B (n=3) vs Ctrl (n=3). Analysi conducted on the total cellular proteome of the recipient cells following imputation with a p-value of <0.05 and an activation z-score of > +/- 2, showing likely decreased expression of (A) EIF3E (dark blue) and increased expression of pathways (B) NR3C1 and SLC40A1 (orange) based on the increased measurements (red) and decreased measurements (light blue) of the mass spectrometry data. Solid lines indicate a direct relationship between proteins, dashed line indicate and indirect relationship. Yellow lines indicate that the pathway activation state does not align with the protein abundance measured in the data.

We identified the top canonical pathways for DA proteins for concentration-A vs control untreated and concentration-B vs control untreated. EV treatment activated pathways involved with cell cycle and cell signalling. Inhibited pathways included protein sorting via vesicles, and fatty acid β-oxidation (**Figure 8 and 9**).

**Figure 8.**
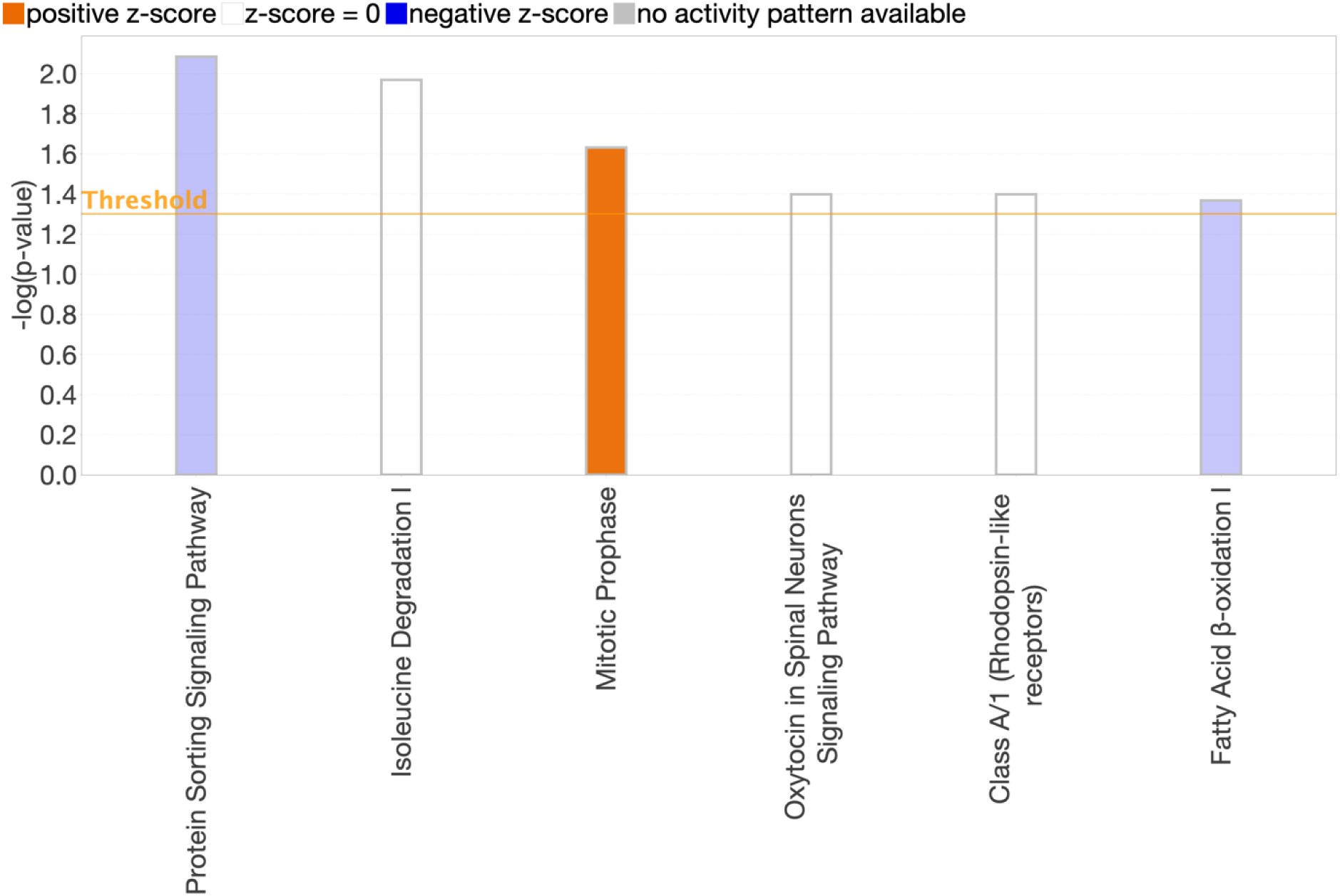
Top canonical pathways identified following treatment with concentration-A v control. Pathway analysis conducted on DE proteins within the total cellular proteome of the recipient cells following imputation (n=3) with thresholds of p-value <0.05 and 2-FC. Pathway in blue are predicted to be inhibited compared to orange predicted to be activated.

**Figure 9.**
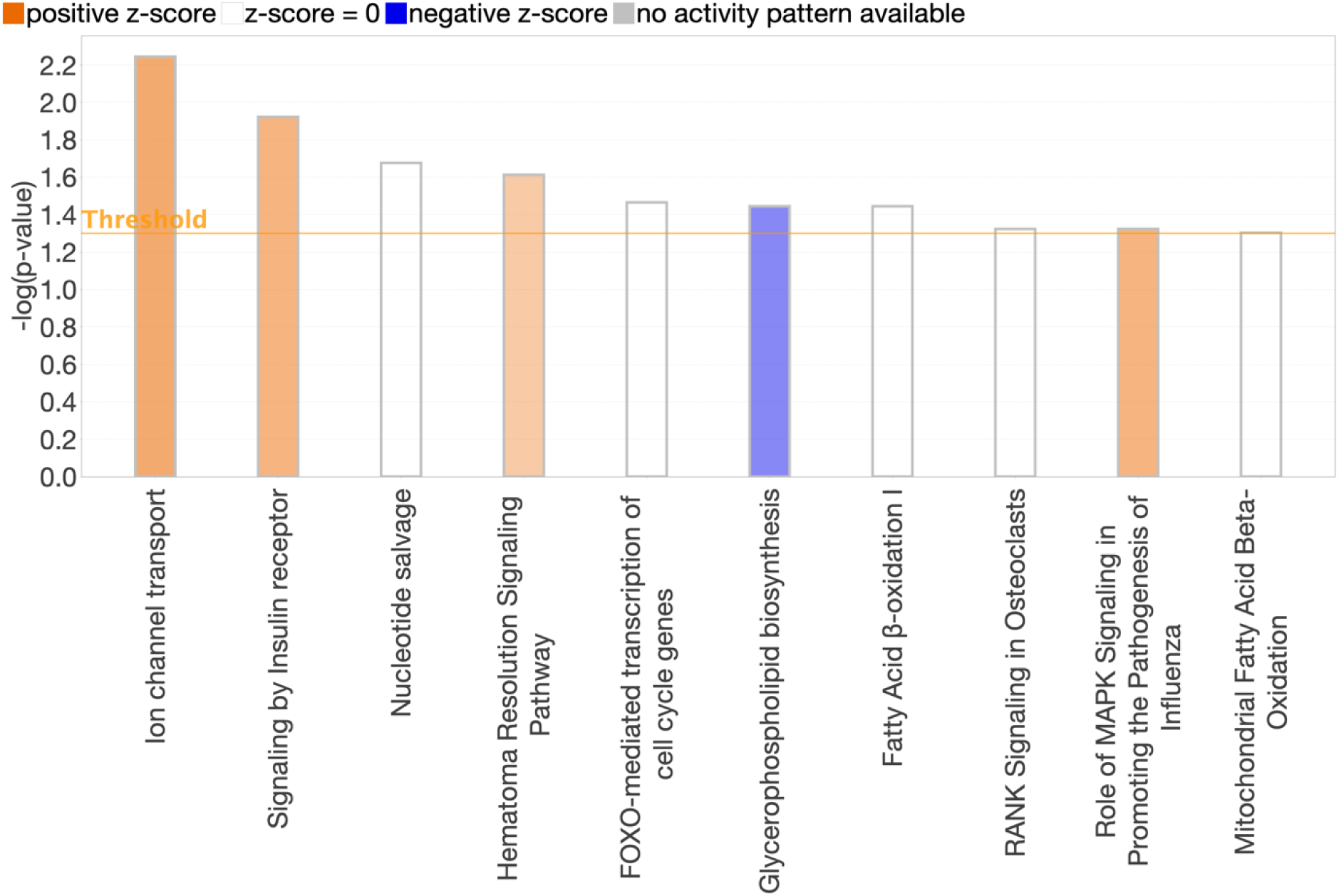
Top canonical pathways identified following treatment with concentration-B v control. Pathway analysis conducted on DE proteins within the total cellular proteome of the recipient cells following imputation (n=3) with thresholds of p-value <0.05 and 2-FC. Pathway in blue are predicted to be inhibited compared to orange predicted to be activated.

## 5. Discussion

We cultured donor cells in heavy labelled media to label the EVs produced. These heavy EVs were then applied onto unlabelled recipient cells for 12 hours to EV uptake at varying concentrations of applied EVs. We detected a heavy signal within the unlabelled recipient cells originating from the heavy donor cell EVs. This is the first study to our knowledge to use mass spectrometry to track protein cargo taken up by recipient cells.

Due to our preliminary results (data not shown) indicating cell stress beginning at 5.5 hours post-treatment, we assessed cell viability following EV treatment for 6 hours in no FCS, or 10% EV-free FCS. However, no effect was observed. It is possible that the EVs were not incubated on the cells for sufficient time to see the effects of these treatments on the cells. Furthermore, cells may have had lower levels of stress which did not affect viability. It is feasible that the cells were stressed but not dying at 6 hours incubation, thus a cell stress assay may provide a better indication of the cell status. EV treatment times can vary between 12 hours to 6 days^26–29^, therefore a longer time-period may have been better.

Combined, FCS and EV tolerance treatment experiments informed our choice to incubate the EVs for heavy-labelling experiments in 10% EV-free FCS. There was no effect of EV concentration on cell viability as measured by visual assessment of confluence. We therefore ranged the EV treatments for the heavy-labelled experiment between 600-6 million EVs/cell. This was to assess whether there was a saturation point of uptake of EVs proxied by an indicated maximal number of heavy transferred proteins or a relationship between the number of transferred proteins and number of EVs applied.

In the heavy-label experiment, the labelling efficiency was 62% following 10 passages of the original donor cells. Passage number was based on previous studies which achieved 91% labelling^30^. The inefficiency could be due to insufficient passage number, or due to labelling reagents being suboptimal. The lack of correlation between the number of EVs applied to the cells for 12 hours and heavy proteins detected within the recipient cells is a significant finding. Our future experiments will use a different batch of heavy label.

We hypothesised that cells would be stressed by physically placing large numbers of EVs onto the cells and this may prevent EVs from being taken up by recipient cells. This could explain why there was no relationship between the number of EVs placed on the cells and the number of transferred heavy-labelled proteins. Interestingly physiological of EVs in human synovial fluid is 1.59×10^10^ particles/mL^31^ which is closest to our concentration-C, compared to concentration-A which was much greater at 1.7×10^12^ particles/mL.

The application of EVs appeared to cause cell stress, supported by the identification of interleukin-37 signalling (an anti-inflammatory pathway unique to treated cell proteins (concentration A-E)). This suggests that applying EVs may have an anti-inflammatory effect. Alternatively, the donor cells (a chondrosarcoma cell line used regularly in cartilage biology research) may have an inflammatory phenotype, and therefore produced the anti-inflammatory protein to reduce inflammation^32^. Alternatively, the recipient cells may have become inflamed by physical application of the EVs and therefore upregulated anti-inflammatory pathways. It is vital that the origin of this inflammation is identified to proceed with this SILAC-based method of transferring proteins. Interleukin-12 signalling was also highlighted in the pathway analysis of the overlapping heavy EV proteomics and heavy recipient cell proteome suggesting pro-inflammatory factors formed by the original donor cells. Interleukin-12 is a pro-inflammatory marker found within the synovial fluid of inflamed osteoarthritic synovial fluid^33^.

We assessed the DA of proteins within the recipient cell proteome following EV treatment. We postulated that varying the amounts of EVs applied to cells would affect the recipient cells, and that by identifying which proteins were altered, we could infer the pathways and mechanisms through which EVs exert their effect in this system. To do this we compared three methods of calculating the number of DA proteins. This was due to the need to deal with missing values in the proteomics data, and as the forced statistical calculations disobeyed assumptions necessary for ANOVA analysis. The most stringent method was imputation in R following ProgenesisQI™ normalisation. Therefore, this method was utilised for DA and the data used for downstream pathway analysis.

We detected heavy-labelled proteins within the recipient cells following treatment of recipient cells with heavy labelled EVs. However further optimisation is necessary for this SILAC-based methodology to be established. Labelling efficiency needs improvement using fresh amino acids and assessment of the optimal number of passages to ensure complete labelling is required. Cell imaging techniques to validate EV uptake should be utilised. The potential inflammatory or anti-inflammatory effect of applying EVs to cells needs investigation to determine whether physically applying high numbers of EVs to cells or transferred protein cargo itself has an effect.

## 6. Conclusion

We found FCS was more influential over the proliferation rate, as a proxy for cell stress, of cells than concentration of EVs applied in our optimisation experiments. However, when we used the proposed optimal concentration of FCS within our heavy experiment, cells still appeared to display indications of stress. We detected heavy proteins within our recipient cells indicating that proteins isolated from the donor cells had been taken up by the recipient cells. This protein cargo overlapped with known EV proteins. There was no relationship between the number of EVs applied to the cells and number of heavy transferred proteins detected. We identified key factors for consideration for follow-up experiments to continue development of this method.

## 7. Acknowledgements

This work was funded by a Horserace Betting Levy Board Project Grant (T15) and British Heart Foundation (SP/F/22/150044) grant.

## Notes

### Competing Interest Statement

The authors have declared no competing interest.

### Summary of Updates

Within this version we have edited the author list with a. change in first author.

## References

1. Peffers MJ, Smagul A, Anderson JR. Proteomic analysis of synovial fluid: current and potential uses to improve clinical outcomes. Expert Rev Proteomics. 2019;16(4):287–302.

2. Rosenthal AK, Gohr CM, Ninomiya J, Wakim BT. Proteomic analysis of articular cartilage vesicles from normal and osteoarthritic cartilage. Arthritis Rheum. 2011;63(2):401–11.

3. Kato T, Miyaki S, Ishitobi H, Nakamura Y, Nakasa T, Lotz MK, et al. Exosomes from IL-1beta stimulated synovial fibroblasts induce osteoarthritic changes in articular chondrocytes. Arthritis Res Ther. 2014;16(4):R163.

4. Yang Z, Feng L, Huang J, Zhang X, Lin W, Wang B, et al. Asiatic acid protects articular cartilage through promoting chondrogenesis and inhibiting inflammation and hypertrophy in osteoarthritis. Eur J Pharmacol. 2021;907:174265.

5. Hoshino A, Kim HS, Bojmar L, Gyan KE, Cioffi M, Hernandez J, et al. Extracellular Vesicle and Particle Biomarkers Define Multiple Human Cancers. Cell. 2020;182(4):1044–61 e18.

6. Anderson JR, Johnson E, Jenkins R, Jacobsen S, Green D, Walters M, et al. Multi-Omic Temporal Landscape of Plasma and Synovial Fluid-Derived Extracellular Vesicles Using an Experimental Model of Equine Osteoarthritis. Int J Mol Sci. 2023;24(19).

7. Es-Haghi M, Godakumara K, Haling A, Lattekivi F, Lavrits A, Viil J, et al. Specific trophoblast transcripts transferred by extracellular vesicles affect gene expression in endometrial epithelial cells and may have a role in embryo-maternal crosstalk. Cell Commun Signal. 2019;17(1):146.

8. Lotvall J, Hill AF, Hochberg F, Buzas EI, Di Vizio D, Gardiner C, et al. Minimal experimental requirements for definition of extracellular vesicles and their functions: a position statement from the International Society for Extracellular Vesicles. J Extracell Vesicles. 2014;3:26913.

9. Thery C, Witwer KW, Aikawa E, Alcaraz MJ, Anderson JD, Andriantsitohaina R, et al. Minimal information for studies of extracellular vesicles 2018 (MISEV2018): a position statement of the International Society for Extracellular Vesicles and update of the MISEV2014 guidelines. J Extracell Vesicles. 2018;7(1):1535750.

10. Witwer KW, Goberdhan DC, O’Driscoll L, Thery C, Welsh JA, Blenkiron C, et al. Updating MISEV: Evolving the minimal requirements for studies of extracellular vesicles. J Extracell Vesicles. 2021;10(14):e12182.

11. Witwer KW, Soekmadji C, Hill AF, Wauben MH, Buzas EI, Di Vizio D, et al. Updating the MISEV minimal requirements for extracellular vesicle studies: building bridges to reproducibility. J Extracell Vesicles. 2017;6(1):1396823.

12. Welsh JA, Goberdhan DCI, O’Driscoll L, Buzas EI, Blenkiron C, Bussolati B, et al. Minimal information for studies of extracellular vesicles (MISEV2023): From basic to advanced approaches. J Extracell Vesicles. 2024;13(2):e12404.

13. Mol EA, Goumans MJ, Doevendans PA, Sluijter JPG, Vader P. Higher functionality of extracellular vesicles isolated using size-exclusion chromatography compared to ultracentrifugation. Nanomedicine. 2017;13(6):2061–5.

14. Nordin JZ, Lee Y, Vader P, Mager I, Johansson HJ, Heusermann W, et al. Ultrafiltration with size-exclusion liquid chromatography for high yield isolation of extracellular vesicles preserving intact biophysical and functional properties. Nanomedicine. 2015;11(4):879–83.

15. Takov K, Yellon DM, Davidson SM. Comparison of small extracellular vesicles isolated from plasma by ultracentrifugation or size-exclusion chromatography: yield, purity and functional potential. J Extracell Vesicles. 2019;8(1):1560809.

16. Finoulst I, Vink P, Rovers E, Pieterse M, Pinkse M, Bos E, et al. Identification of low abundant secreted proteins and peptides from primary culture supernatants of human T-cells. J Proteomics. 2011;75(1):23–33.

17. Munoz J, Heck AJ. Quantitative proteome and phosphoproteome analysis of human pluripotent stem cells. Methods Mol Biol. 2011;767:297–312.

18. Debaisieux S, Encheva V, Chakravarty P, Snijders AP, Schiavo G. Analysis of Signaling Endosome Composition and Dynamics Using SILAC in Embryonic Stem Cell-Derived Neurons. Mol Cell Proteomics. 2016;15(2):542–57.

19. Scheffer LL, Sreetama SC, Sharma N, Medikayala S, Brown KJ, Defour A, et al. Mechanism of Ca(2)(+)-triggered ESCRT assembly and regulation of cell membrane repair. Nat Commun. 2014;5:5646.

20. Johnson BB, Cosson MV, Tsansizi LI, Holmes TL, Gilmore T, Hampton K, et al. Perlecan (HSPG2) promotes structural, contractile, and metabolic development of human cardiomyocytes. Cell Rep. 2024;43(1):113668.

21. Cox J, Matic I, Hilger M, Nagaraj N, Selbach M, Olsen JV, et al. A practical guide to the MaxQuant computational platform for SILAC-based quantitative proteomics. Nat Protoc. 2009;4(5):698–705.

22. Pathan M, Keerthikumar S, Ang CS, Gangoda L, Quek CY, Williamson NA, et al. FunRich: An open access standalone functional enrichment and interaction network analysis tool. Proteomics. 2015;15(15):2597–601.

23. Pathan M, Keerthikumar S, Chisanga D, Alessandro R, Ang CS, Askenase P, et al. A novel community driven software for functional enrichment analysis of extracellular vesicles data. J Extracell Vesicles. 2017;6(1):1321455.

24. Kalra H, Simpson RJ, Ji H, Aikawa E, Altevogt P, Askenase P, et al. Vesiclepedia: a compendium for extracellular vesicles with continuous community annotation. PLoS Biol. 2012;10(12):e1001450.

25. Mellman I, Nelson WJ. Coordinated protein sorting, targeting and distribution in polarized cells. Nat Rev Mol Cell Biol. 2008;9(11):833–45.

26. Jeon OH, Wilson DR, Clement CC, Rathod S, Cherry C, Powell B, et al. Senescence cell-associated extracellular vesicles serve as osteoarthritis disease and therapeutic markers. JCI Insight. 2019;4(7).

27. Bhome R, Emaduddin M, James V, House LM, Thirdborough SM, Mellone M, et al. Epithelial to mesenchymal transition influences fibroblast phenotype in colorectal cancer by altering miR-200 levels in extracellular vesicles. J Extracell Vesicles. 2022;11(5):e12226.

28. Liu W, Liu A, Li X, Sun Z, Sun Z, Liu Y, et al. Dual-engineered cartilage-targeting extracellular vesicles derived from mesenchymal stem cells enhance osteoarthritis treatment via miR-223/NLRP3/pyroptosis axis: Toward a precision therapy. Bioact Mater. 2023;30:169–83.

29. Sun Y, Zhao J, Wu Q, Zhang Y, You Y, Jiang W, et al. Chondrogenic primed extracellular vesicles activate miR-455/SOX11/FOXO axis for cartilage regeneration and osteoarthritis treatment. NPJ Regen Med. 2022;7(1):53

30. Li N, Li J, Desiderio DM, Zhan X. SILAC quantitative proteomics analysis of ivermectin-related proteomic profiling and molecular network alterations in human ovarian cancer cells. J Mass Spectrom. 2021;56(1):e4659.

31. Withrow J, Murphy C, Dukes A, Fulzele S, Liu Y, Hunter M, et al., editors. Synovial fluid exosomal MicroRNA profiling of osteoarthritis patients and identification of synoviocyte-chondrocyte communication pathway. ORS 2016 Annual Meeting; 2016.

32. Nold MF, Nold-Petry CA, Zepp JA, Palmer BE, Bufler P, Dinarello CA. IL-37 is a fundamental inhibitor of innate immunity. Nat Immunol. 2010;11(11):1014–22.

33. Moradi B, Rosshirt N, Tripel E, Kirsch J, Barie A, Zeifang F, et al. Unicompartmental and bicompartmental knee osteoarthritis show different patterns of mononuclear cell infiltration and cytokine release in the affected joints. Clin Exp Immunol. 2015;180(1):143–54.

